## Supplementary information for "Disentangling acute motor deficits and adaptive responses evoked by the loss of cerebellar output"

**Table S1: Mean success rate across sessions per monkey.**

| Monkey | Success rate (Control) | Success rate (Cerebellar block) |
|---|---|---|
| Monkey S | 84.9% CI [82.2, 87.5] | 66.8% CI [63.4, 70.3] |
| Monkey C | 89.9% CI [88.8, 90.9] | 82.3% CI [79.8, 84.7] |
| Monkey M | 80.4% CI [79.1, 81.6] | 66.1% CI [63.4, 68.7] |
| Monkey P | 90.4% CI [89.3, 91.5] | 82.4% CI [80.0, 84.6] |

**Table S2: ANOVA marginal tests for the effect of target direction on the change in peak hand velocity due to cerebellar block relative to control. (DF: degrees of freedom)**

| **Model: Peak velocity change ~ Target + (1 + Target \| Subject)** | | | | |
|---|---|---|---|---|
| **Term** | **F-Statistic** | **DF1** | **DF2** | **p-value** |
| Intercept | 6.66 | 1 | 774 | 0.01 |
| Target | 4.89 | 7 | 774 | < 0.001 |

**Description:** Movements exhibited a significant reduction in peak hand velocity during the cerebellar block in a target-dependent manner. The change in peak hand velocity was modeled using a linear mixed-effects model, with target as a fixed effect and random intercepts and slopes for target within each subject (i.e. monkey). For each session, the target-wise change in the median peak hand velocity during the cerebellar block trials was computed relative to that of control trials. The input to the model was the target-wise values computed from all sessions pooled across all four monkeys. The significant effect of target direction on the change in peak velocity can be interpreted as analogous to the interaction between cerebellar block and target direction on the actual peak velocities.

**Table S3a: ANOVA marginal tests for the effect of target group (targets 1-4 vs. targets 5-8) on peak hand velocity during control. (DF: degrees of freedom)**

| Model: Peak velocity ~ Target group + (1 + Target group \| Subject) | | | | |
|---|---|---|---|---|
| **Term** | **F-Statistic** | **DF1** | **DF2** | **p-value** |
| Intercept | 114.80 | 1 | 774 | < 0.001 |
| Target group | 0.02 | 7 | 774 | 0.889 |

**Description:** Movements exhibited no significant difference in peak hand velocity during the outward reaching (targets 1-4) vs. retrieval (targets 5-8) movements during the control condition. The peak hand velocity was modeled using a linear mixed-effects model, with the target group (T1-4 vs. T5-8) as a fixed effect and random intercepts and slopes for the target group within each subject (i.e. monkey). The median peak hand velocity during the control trials was computed for each session. The input to the model was the target group-wise values computed from all sessions pooled across all four monkeys.

**Table S3b: Mean peak hand velocity for outward reaching (targets 1-4) vs. inward reaching (targets 5-8) movements during control across all sessions per monkey.** For each session, the median peak hand velocity was computed across all the control trials per target group. Then, the mean and confidence intervals of the mean were computed from the per-session data for each monkey.

| Monkey | Targets 1-4 (mean ± CI, cm/s) | Targets 5-8 (mean ± CI, cm/s) |
|---|---|---|
| Monkey S | 18.2 [17.3, 19.0] | 18.4 [17.5, 19.1] |
| Monkey C | 12.5 [11.4, 13.0] | 11.5 [10.9, 12.8] |
| Monkey M | 14.6 [14.3, 15.1] | 15.1 [14.8 15.5] |
| Monkey P | 11.2 [10.8, 11.6] | 12.3 [11.9, 12.7] |

**Table S4: ANOVA marginal tests for the effect of target direction on the change in shoulder muscle torque impulse due to cerebellar block relative to control. (DF: degrees of freedom)**

| **Model: Shoulder muscle torque change ~ Target + (1 + Target \| Subject)** | | | | |
|---|---|---|---|---|
| **Term** | **F-Statistic** | **DF1** | **DF2** | **p-value** |
| Intercept | 7.22 | 1 | 774 | 0.01 |
| Target | 12.55 | 7 | 774 | < 0.001 |

**Description:** Movements exhibited a significant reduction in shoulder muscle torque impulse during the cerebellar block in a target-dependent manner. The torque impulse was computed by integrating the torque profile during the positive acceleration phase of the movement. The change in muscle torque impulse was modeled using a linear mixed-effects model, with target as a fixed effect and random intercepts and slopes for target within each subject (i.e. monkey). For each session, the target-wise change in the median muscle torque impulse during the cerebellar block trials was computed relative to that of control trials. The input to the model was the target-wise values computed from all sessions pooled across all four monkeys. The significant effect of target direction on the change in muscle torque impulse can be interpreted as analogous to the interaction between cerebellar block and target direction on the actual peak velocities.

**Table S5: ANOVA marginal tests for the effect of target direction on the change in elbow muscle torque impulse due to cerebellar block relative to control. (DF: degrees of freedom)**

| Model: Elbow muscle torque change ~ Target + (1 + Target \| Subject) | | | | |
|---|---|---|---|---|
| **Term** | **F-Statistic** | **DF1** | **DF2** | **p-value** |
| Intercept | 0.46 | 1 | 774 | 0.50 |
| Target | 0.66 | 7 | 774 | 0.70 |

**Description:** Movements exhibited a significant reduction in elbow muscle torque impulse during the cerebellar block in a target-dependent manner. The torque impulse was computed by integrating the torque profile during the positive acceleration phase of the movement. The change in muscle torque impulse was modeled using a linear mixed-effects model, with target as a fixed effect and random intercepts and slopes for target within each subject (i.e. monkey). For each session, the target-wise change in the median muscle torque impulse during the cerebellar block trials was computed relative to that of control trials. The input to the model was the target-wise values computed from all sessions pooled across all four monkeys.

**Table S6: ANOVA marginal tests for the effect of trial sequence and trial type (control/cerebellar block) on the peak hand velocity relative to 1st 2 trials in control for movements to target 1. (DF: degrees of freedom)**

| **Model: Peak velocity (%) ~ Trial type x Trial sequence + (1 + Trial type x Trial sequence \| Subject)** | | | | |
|---|---|---|---|---|
| **Term** | **F-Statistic** | **DF1** | **DF2** | **p-value** |
| Intercept | 7472.51 | 1 | 293 | < 0.001 |
| Trial type | 8.35 | 1 | 293 | 0.004 |
| Trial sequence | 0.03 | 1 | 293 | 0.957 |
| Trial type : Trial sequence | 0.16 | 1 | 293 | 0.693 |

**Description:** The evolution of peak hand velocities during control vs. cerebellar block was analyzed by preserving the order of presentation of the target in each block of trials (i.e., trial sequence). For each monkey, the peak hand velocities were normalized to the median peak velocity of the early trials 1-2 in the control blocks. The normalized peak velocities were then modeled using a linear mixed-effects model, with trial type (control/cerebellar block) and trial sequence (1-20) as fixed effects and random intercepts and slopes for trial type and trial sequence within each subject (i.e. monkey).

**Table S7: ANOVA marginal tests for the effect of trial sequence and trial type (control/cerebellar block) on the peak hand velocity relative to 1st 2 trials in control for movements to target 2-4. (DF: degrees of freedom)**

| Model: Peak velocity (%) ~ Trial type x Trial sequence + (1 + Trial type x Trial sequence \| Subject) | | | | |
|---|---|---|---|---|
| **Term** | **F-Statistic** | **DF1** | **DF2** | **p-value** |
| Intercept | 23585.00 | 1 | 1178 | < 0.001 |
| Trial type | 18.29 | 1 | 1178 | < 0.001 |
| Trial sequence | 5.48 | 1 | 1178 | 0.02 |
| Trial type : Trial sequence | 7.30 | 1 | 1178 | 0.007 |

**Description:** The evolution of peak hand velocities during control vs. cerebellar block was analyzed by preserving the order of presentation of the target in each block of trials (i.e., trial sequence). For each monkey, the peak hand velocities were normalized to the median peak velocity of the early trials 1-2 in the control blocks. The normalized peak velocities were then modeled using a linear mixed-effects model, with trial type (control/cerebellar block) and trial sequence (1-20) as fixed effects and random intercepts and slopes for trial type and trial sequence within each subject (i.e. monkey).

**Table S8: ANOVA marginal tests for the effect of trial sequence and trial type (control/cerebellar block) on movement decomposition relative to 1st 2 trials in control for movements to targets 2-4. (DF: degrees of freedom)**

| **Model: Decomposition index (%) ~ Trial type x Trial sequence + (1 + Trial type x Trial sequence \| Subject)** | | | | |
|---|---|---|---|---|
| **Term** | **F-Statistic** | **DF1** | **DF2** | **p-value** |
| Intercept | 884.41 | 1 | 1021 | < 0.001 |
| Trial type | 52.97 | 1 | 1021 | < 0.001 |
| Trial sequence | 2.08 | 1 | 1021 | 0.149 |
| Trial type : Trial sequence | 0.63 | 1 | 1021 | 0.426 |

**Description:** The evolution of decomposition index (measured as the fraction of time during a movement when either, but not both, of the shoulder or the elbow joint velocity was less than 20°/s) during control vs. cerebellar block was analyzed by preserving the order of presentation of the target in each block of trials (i.e., trial sequence). For each monkey, the decomposition indices were normalized to the median decomposition index of the early trials 1-2 in the control blocks. The normalized decomposition indices were then modeled using a linear mixed-effects model, with trial type (control/cerebellar block) and trial sequence (1-20) as fixed effects and random intercepts and slopes for trial type and trial sequence within each subject (i.e. monkey).

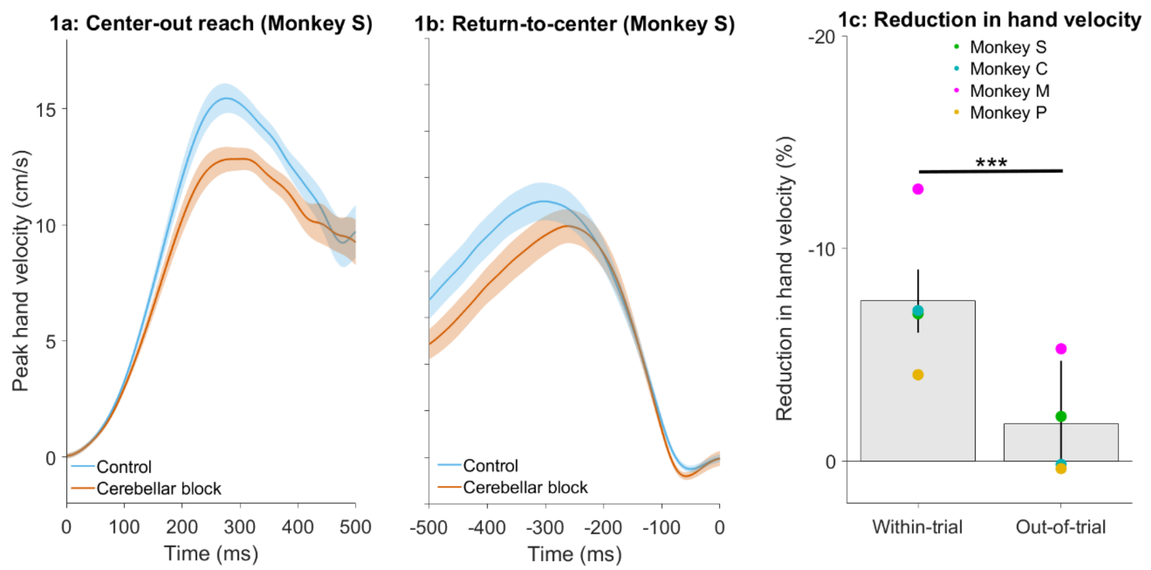


**Figure S1: Effect of cerebellar block on peak hand velocity during within-trial vs. inter-trial movements.** (a) Movement-onset aligned hand velocity profiles of target-directed reaching movements performed by monkey S. We took the median velocity profile per target per session then averaged across all targets to generate one velocity profile per session. The depicted data is the mean ± 95% confidence intervals across all sessions. Movement onset was defined as the first time point when the hand velocity exceeded 5% of the peak. To facilitate visualization, we subtracted the individual baseline (mean of the 1st 10 ms from movement onset) from the plotted traces. (b) End-of-movement aligned hand velocity profiles of return-to-center movements performed by monkey S. We took the median velocity profiles per target per session and then averaged across all targets to generate one velocity profile per session. The depicted data is the mean ± 95% confidence intervals across all sessions. Movement end was defined as the last time point when the hand velocity exceeded 5% of the peak. To facilitate visualization, we subtracted the individual baseline (mean of the last 10 ms before the end of movement) from the plotted traces. (c) Effect of the cerebellar block on peak hand velocity for within-trial target-directed movements vs. out-of-trial return-to-center movements. For each session, the target-wise reduction in the median peak hand velocity during the cerebellar block trials was computed relative to that of control trials. The depicted values are the means ± 95% confidence intervals across all sessions pooled from all four monkeys. The means of individual monkeys are overlaid. Statistical significance is denoted as follows: $p \geq 0.05$NS, $p < 0.05$*, $p < 0.01$**, $p < 0.001$***.

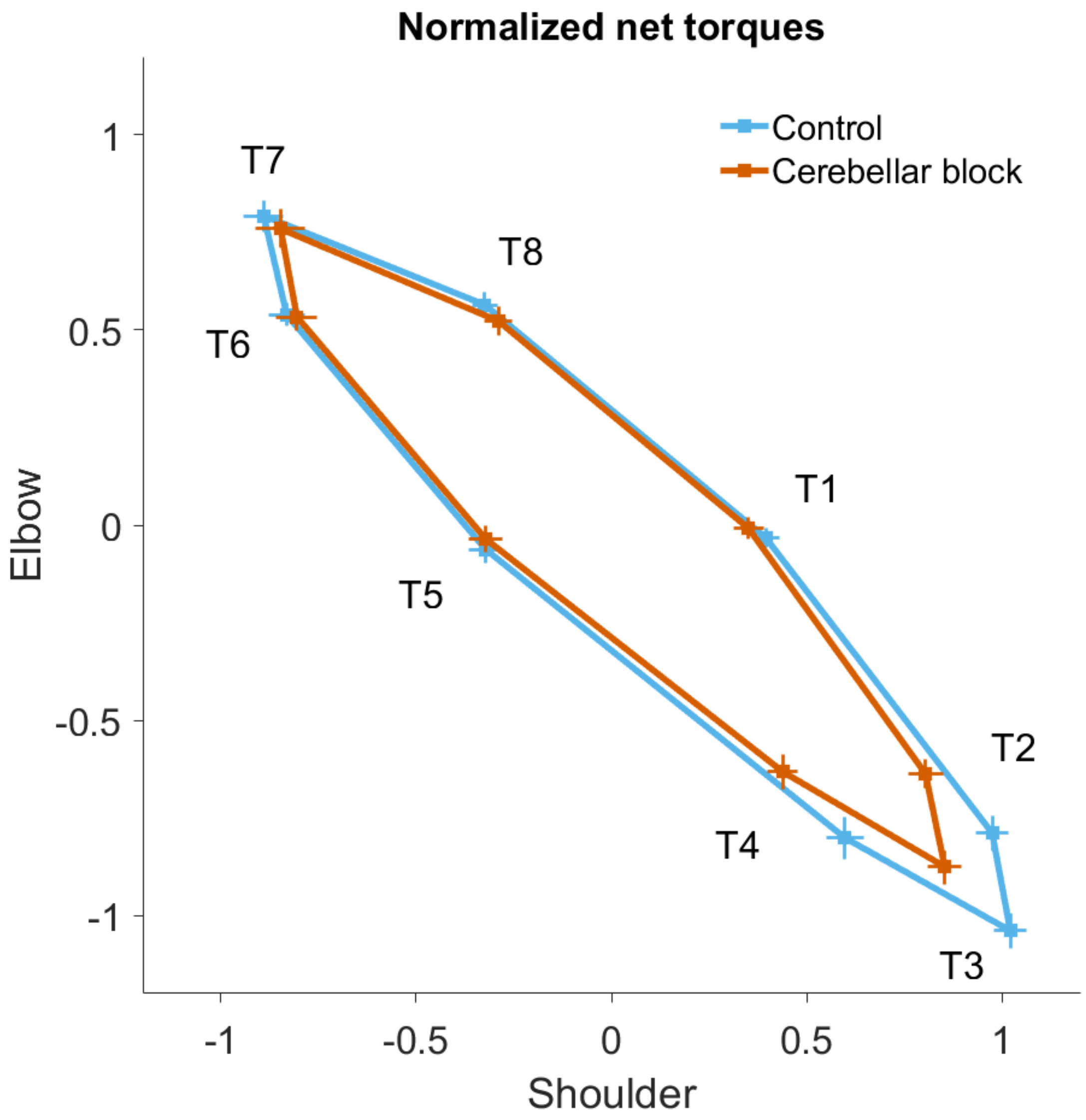


**Supplementary Figure S2:** Normalized net torque impulse at the shoulder vs. elbow per target during control and cerebellar block conditions. For each session, we computed the target-wise median net torque impulse during the acceleration phase of the movement across the control and cerebellar block trials. They were then normalized by the maximum absolute torque impulse across all the targets. The depicted values are the means across all the sessions pooled from the data of all four monkeys. The 95% confidence interval of the means is indicated by the horizontal and vertical bars for the shoulder and elbow joints respectively.

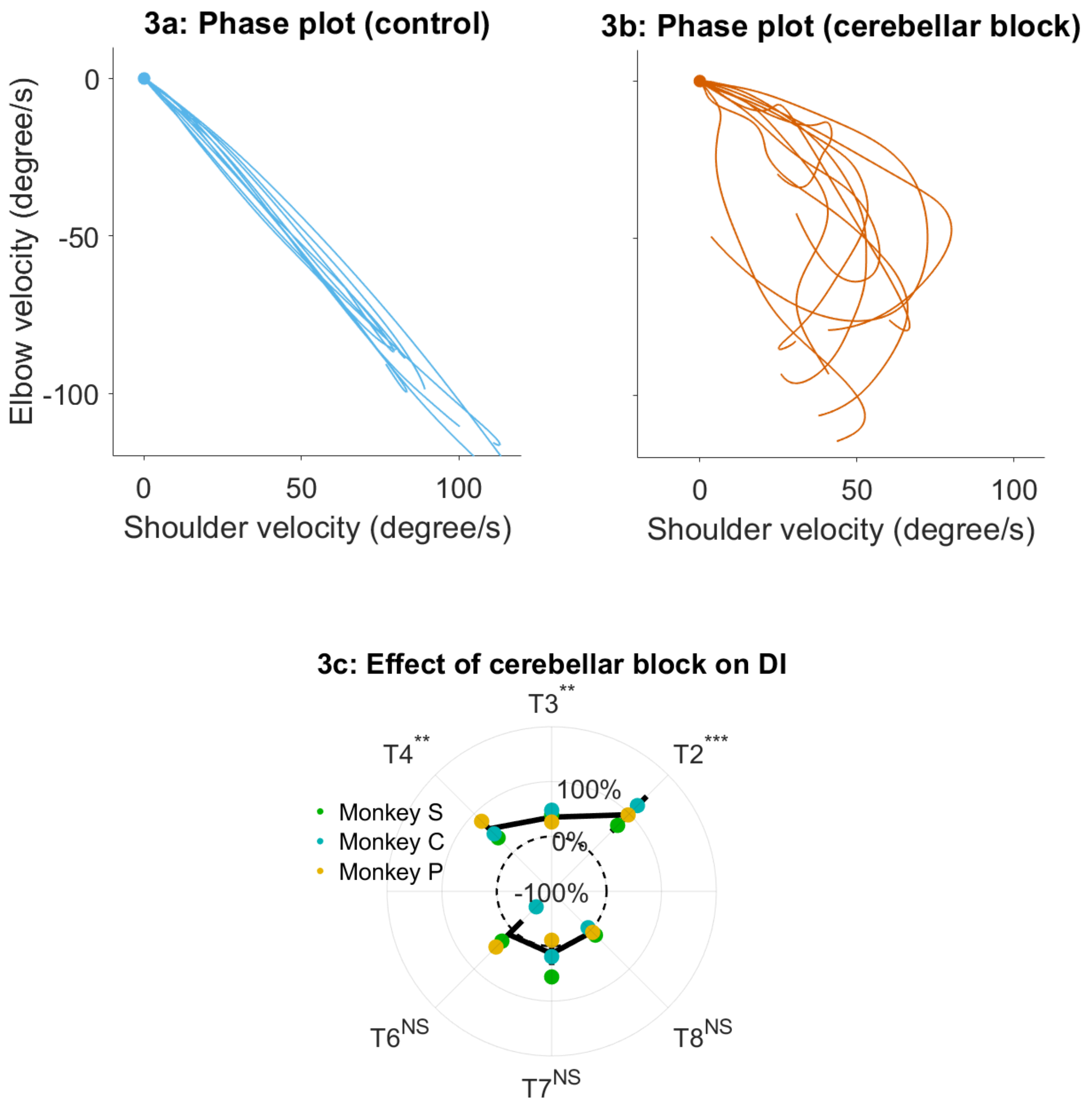


**Supplementary Figure S3a-c: Effect of the cerebellar block on the decomposition of movements.** (a) Phase plot of elbow versus shoulder angular velocity for 10 representative zero-centered control trials (Monkey S, target 2). (b) Same as (a), but during cerebellar block. Contrasting the two plots reveals synchronous joint movements under control conditions versus asynchronous joint movements during the cerebellar block, indicating movement decomposition in the latter. (c) Change in decomposition index (i.e., the proportion of the movement time during which the movement was decomposed) for movements to each target during velocity-matched cerebellar block trials (The change in decomposition was computed per session per target as: [median decomposition index of velocity matched cerebellar block trials – median decomposition of velocity matched control trials]/median decomposition of all control trials x 100). The depicted values are the mean ± 95% confidence intervals across all sessions pooled from the individual monkeys. The individual means of each monkey are overlaid. We removed monkey M from this analysis to improve power for the statistical tests after correction for multi-comparisons since Monkey M exhibited overall low decomposition. Targets 1 and 5 were excluded from this analysis since the elbow joint is mostly stationary for movements to these targets (i.e., the net torque at the elbow is very low as shown in Supplementary figure S2). Statistical significance is denoted as follows: $p \geq 0.05$NS, $p < 0.05$*, $p < 0.01$**, $p < 0.001$*** [T1-8: Targets 1-8]

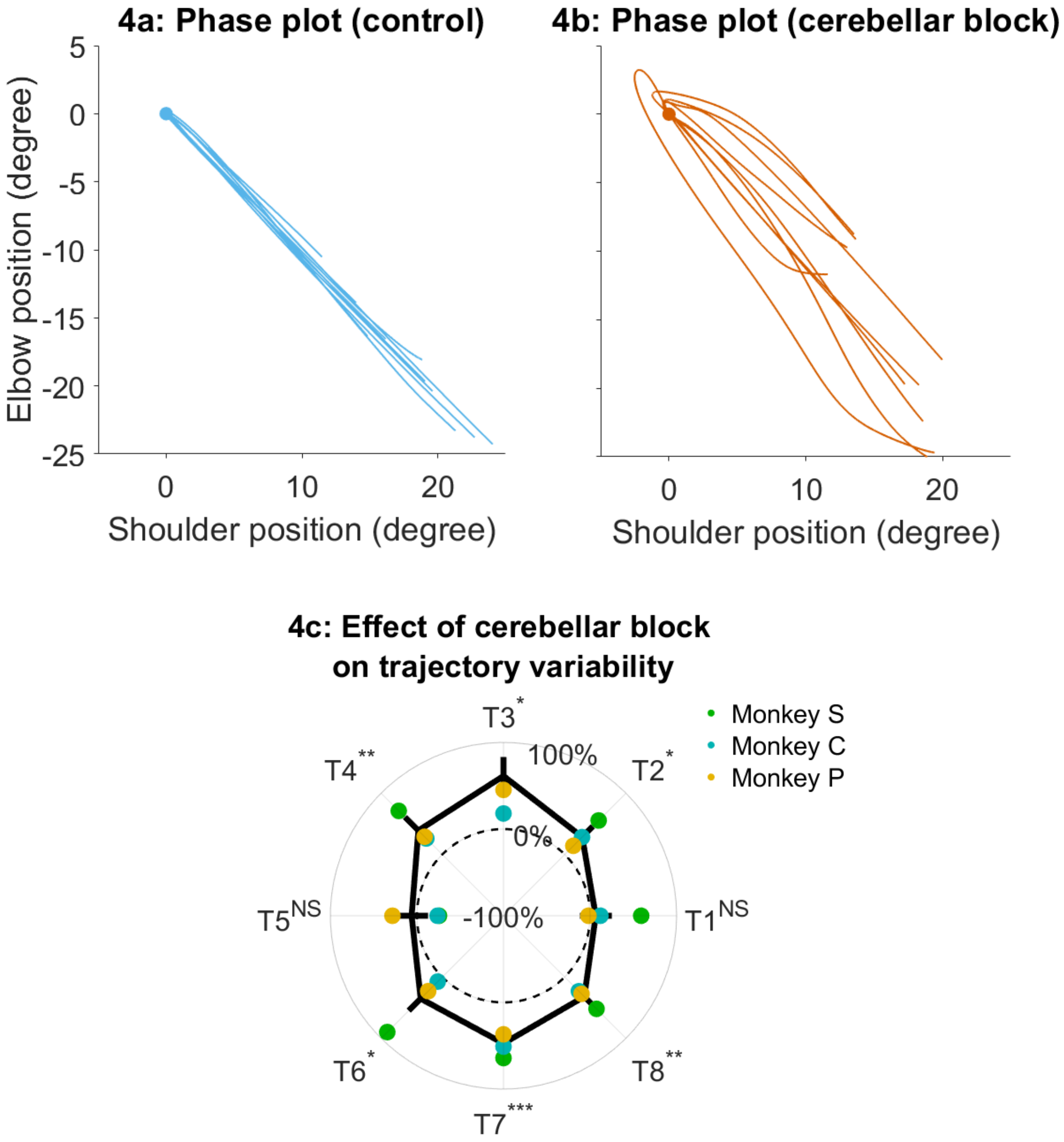


**Supplementary Figure S4a-c: Effect of the cerebellar block on the trajectory variability.** (a) Phase plot of elbow versus shoulder angular position for 10 representative zero-centered control trials (Monkey S, target 2). (b) Same as (a), but during cerebellar block. Contrasting the two plots reveals the increased inter-trial variability in the shoulder-elbow coordination during cerebellar block relative to control which led to increased trajectory variability. (c) Change in inter-trial trajectory variability for movements to each target during velocity-matched cerebellar block trials. The trajectory variability was measured as the standard deviation of the maximum perpendicular distance of the trajectories from the Y-axis after transforming them as in Figure 5d of the main text. The change in trajectory variability for the cerebellar block trials was computed relative to the control trials per session per target. The depicted values are the mean ± 95% confidence intervals across all sessions pooled from the individual monkeys. The individual means of each monkey are overlaid. We removed monkey M from this analysis to improve power for the statistical tests after correction for multi-comparisons since Monkey M exhibited overall low trajectory variability. Statistical significance is denoted as follows: $p \geq 0.05$NS, $p < 0.05$*, $p < 0.01$**, $p < 0.001$*** [T1-8: Targets 1-8]

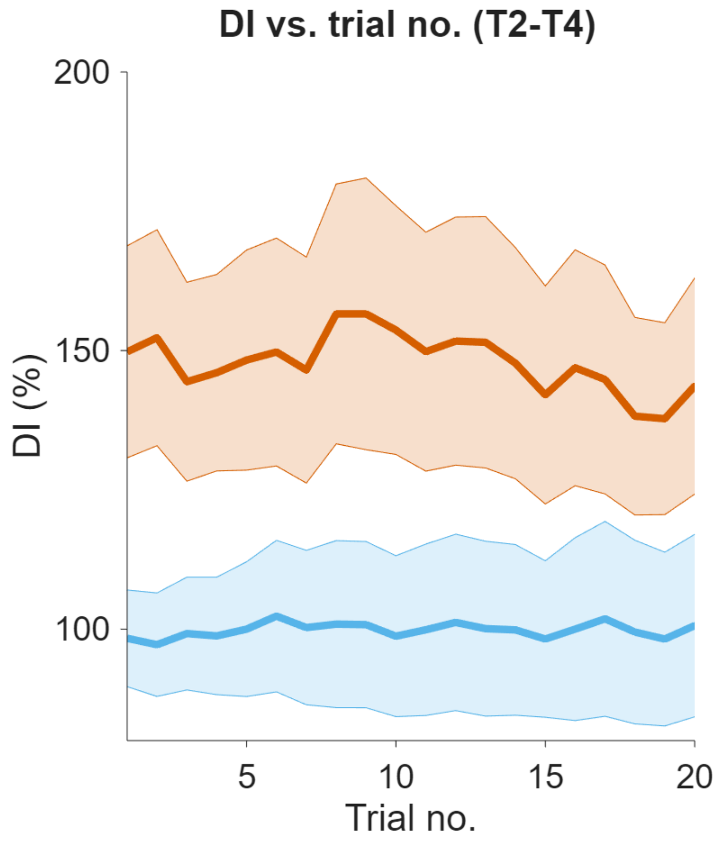


**Supplementary Figure S5: Effect of the cerebellar block on decomposition index across successive trials for movements to targets 2-4.** Decomposition index was computed as the fraction of time during a movement when either (but not both) of the shoulder or the elbow joint velocity was less than 20°/s. The decomposition indices for movements to a particular target were extracted from blocks of control/cerebellar block trials while preserving the order in which the target was presented. For each block of trials, decomposition indices for movements to a particular target was extracted while retaining the trial numbers in which it was presented in the block. This enabled us to examine the evolution of the decomposition index across the sequence of presentation of the target during control vs. cerebellar block. The figure depicts the mean ± 95% confidence intervals of the decomposition index for movements to targets 2-4 across all the trial blocks pooled from all four monkeys. For each monkey, the decomposition indices were normalized (%) to the median decomposition index of the early trials 1-2 from all the control blocks. [DI = Decomposition index, T2-4: Targets 2-4]
